# The Developmental Profile of Visual Cortex Astrocytes

**DOI:** 10.1101/2022.09.27.509759

**Authors:** Airi Watanabe, Connie Guo, Per Jesper Sjöström

## Abstract

Though once thought of as passive support cells for neurons, it is now clear that astrocytes signal via calcium (Ca^2+^) to trigger gliotransmission that impacts surrounding neurons and synapses. We therefore investigated how astrocytes in layer-5 (L5) mouse visual cortex mature electrophysiologically and morphologically over postnatal days (P) 3 – 30 and how these relate to changes in spontaneous Ca^2+^ events. Across this age range, resting membrane potential increased, and input resistance decreased. Astrocytes also revealed different membrane responses to voltage steps: notably, membrane responses became more passive with age. Two-photon (2p) imaging of dye-loaded patched cells and confocal imaging revealed that gap-junction coupling increased starting around P7. Morphological reconstructions revealed increased branch density but also shorter branches after P20, suggesting that astrocyte branches may get pruned as dense tiling is established. Finally, we visualized spontaneous Ca^2+^ transients with 2p microscopy and found that Ca^2+^ events decorrelated, became more frequent as well as briefer with age. As astrocytes mature, spontaneous Ca^2+^ activity thus changes from relatively cell-wide, synchronous waves to local transients. Several astrocyte properties were stably mature from ~P15 onwards, although morphology continued to develop further. Our findings provide a descriptive foundation of astrocyte maturation, useful for the study of astrocytic impact on visual cortex critical period plasticity.

## INTRODUCTION

Historically, astrocytes have been considered structural support cells in the central nervous system, meaning they hold neuronal components together, maintain molecular homeostasis, and control the blood-brain-barrier (Verkhratsky and Nedergaard, 2018). One reason that astrocytes were chiefly thought of as supportive is that their excitability is poor compared to neurons, e.g., astrocytes cannot propagate action potentials. However, in the early 1990s, it was discovered that these glia cells do in fact exhibit active responses, although via Ca^2+^ signals (Charles et al., 1991; Cornell-Bell et al., 1990). Soon after this discovery, it was established that, following cytosolic Ca^2+^ elevation, astrocytes respond to neurotransmitters by releasing gliotransmitters (Nedergaard, 1994; Parpura et al., 1994). Moreover, stimulation of a single astrocyte can trigger a wave of intracellular Ca^2+^ that propagates from astrocyte to astrocyte via gap junctions — channels specialized for cell-to-cell communication (Giaume et al., 2010; Nedergaard, 1994).

Astrocytes have since been implicated in several central nervous system functions, including synaptic plasticity (Ribot et al., 2021) and neurotransmitter buffering (Weiss et al., 2019). Importantly, a single mature astrocyte contacts over 100,000 synapses with each astrocyte occupying tiles, i.e., exclusive, non-overlapping territories (Bushong et al., 2002). Therefore, they are in strategic positions to modulate synapses within tiles. On the other hand, gap junctions allow Ca^2+^ signals to propagate in the interconnected astrocyte syncytium network. Gap-junction coupling thus expands the modulation range of astrocytes beyond the tile (Pannasch and Rouach, 2013).

Ca^2+^ signaling in astrocytes occurs either spontaneously or in response to neurotransmitter stimulation (Aguado et al., 2002; Perea and Araque, 2005a). These Ca^2+^ signals — which are relevant for basal synaptic function (Di Castro et al., 2011) — originate in microdomains of astrocyte processes and propagate to other regions of the cell (Grosche et al., 1999; Perea and Araque, 2005b) as well as to neighboring astrocytes (Fiacco and McCarthy, 2004). This astrocyte Ca^2+^ activity triggers the release of gliotransmitters, such as D-serine, ATP, and glutamate (Volterra and Meldolesi, 2005). For example, astrocyte Ca^2+^ signaling actively participates in the induction of neocortical spike-timing-dependent plasticity (Min and Nevian, 2012). A key remaining question, however, is how the morphological development of astrocytes is coordinated with the maturation of intracellular Ca^2+^ signaling, which has implications for astrocytic control of neocortical plasticity (Min and Nevian, 2012).

We therefore explored how electrophysiological and morphological properties of layer 5 (L5) visual cortex astrocytes mature with age. We found that, over development, astrocytes elaborated denser arborizations, spontaneous Ca^2+^ activity in individual astrocytes decorrelated, and Ca^2+^ events increased in frequency as well as decreased in duration. Several astrocyte properties seemed to stabilize by P15, including gap-junction coupling, membrane biophysics, and Ca^2+^ signaling. Astrocyte morphology, however, matured well after P20, by becoming denser, more compact, and more symmetric.

## METHODS

### Animals and ethics statement

The animal study was reviewed and approved by the Montreal General Hospital Facility Animal Care Committee (The MGH FACC) and adhered to the guidelines of the Canadian Council on Animal Care (CCAC). Female and male C57BL/6J mice from P3 – P30 were anesthetized with isoflurane and sacrificed by decapitation once the hind-limb withdrawal reflex was lost.

### Acute slice electrophysiology

After decapitation, the brain was removed and placed in ice-cold (~4°C) artificial cerebrospinal fluid (ACSF), containing in mM: 125 NaCl, 2.5 KCl, 1 MgCl_2_, 1.25 NaH_2_PO_4_, 2 CaCl_2_, 26 NaHCO_3_ and 25 glucose, bubbled with 95% O_2_/5% CO_2_ (carbogen). Osmolality of the ACSF was adjusted to ~338 mOsm with glucose. Oblique coronal 300-μm-thick acute brain slices were prepared using a Campden Instruments 5000 mz-2 vibratome (Lafayette Instrument, Lafayette, IN, USA). Brain slices were kept at ~33°C in oxygenated ACSF for ~30 min and then allowed to cool at room temperature for at least one hour after slicing before patching.

Glass pipettes (3 – 7 MΩ) were used for whole-cell recordings and were filled with internal solution (in mM: KCl, 5; K-Gluconate, 115; HEPES, 10; Mg-ATP, 4; Na-GTP, 0.3; Na-Phosphocreatine, 10; Biocytin, 0.1% w/v; adjusted with KOH to pH 7.2 and with sucrose to ~310 mOsm, and supplemented with Alexa Fluor 594 or 488, 20 - 80 μM). We carried out electrophysiology experiments with ACSF heated to 32-34°C with a resistive inline heater (Scientifica Ltd, UK), with temperature continuously recorded and verified offline.

BVC-700A (Dagan Corporation, Minneapolis, MN) amplifiers were used to obtain whole-cell recordings. Amplified signals were filtered at 5 kHz and sampled at 40 kHz using PCI-6229 boards (NI, Austin, TX, USA) with MultiPatch custom software (available at https://github.com/pj-sjostrom/MultiPatch.git) running in Igor Pro 8 or 9 (WaveMetrics Inc., Lake Oswego, OR, USA).

Astrocytes were selectively stained by incubating slices in 1 or 5 μM sulforhodamine 101 (SR101) solution (Nimmerjahn and Helmchen, 2012; Nimmerjahn et al., 2004) made up in ACSF for 5 minutes at room temperature while bubbling with carbogen. SR101-positive astrocytes were then visualized with two-photon (2p) microscopy at 820 nm or 930 nm and targeted for whole-cell patching. After patching in current clamp, cell identity was confirmed by checking for a low resting membrane potential (V_m_; ~−70 mV), low input resistance (R_input_; <100 MΩ), and by injecting currents from −0.3 to 0.7 nA at 0.2 nA increments to verify that they do not exhibit action potentials. Resting Vm and R_input_ of each cell were measured from a 250-ms-long 25-pA hyperpolarizing test pulse at the beginning of each current step and taken as an average from the six waves. Additionally, bushy morphology was confirmed post-hoc with 2p imaging. Recordings were not corrected for liquid junction potential (~10 mV) or for series resistance.

To generate IV curves, astrocytes were voltage clamped at −80 mV and voltage steps from +60 mV to −160 mV at 20 mV decrements were applied for 500 ms. To quantify the passivity of the astrocyte, current readings were taken at 10 ms after the beginning of the voltage step (“Early”) and 10 ms before the end of the voltage step (“Late”) and the difference between the slope of the last 5 Late and last 5 Early readings were taken as the change in conductance (ΔConductance) across the voltage step over age.

### Biocytin staining and confocal imaging

Astrocytes were recorded in whole-cell configuration for at least 15 minutes to allow for biocytin diffusion. To ensure re-sealing of cell membrane during pipette removal, cells were held at a depolarizing potential (−40 to −30 mV) while the pipette was slowly removed along the diagonal axis. The acute slice was fixed in 4% paraformaldehyde (PFA) overnight at 4°C and then transferred to 0.1 M phosphate buffered saline (PBS) for up to one week before staining.

Acute slices were washed four times in 10 mM Tris-buffered saline (TBS) solution with 0.3% Triton-X for 10 minutes each. Slices were blocked with 10 mM TBS with 0.3% Triton-X and 10% normal donkey serum (NDS; 017-000-121 Jackson ImmunoResearch, West Grove, PA, USA) for 1 hour. Alexa Fluor 647-conjugated streptavidin (S32357 ThermoFisher Scientific, Waltham, MA, USA) at 1:200 dilution in 0.01 M TBS with 0.3% Triton-X and 1% NDS was used to bind to biocytin overnight at 4°C. Slices were then washed four times in 10 mM TBS solution for 10 minutes each. The slices were mounted on glass slides with ProLong Gold Antifade Mountant (ThermoFisher Scientific, Waltham, MA, USA) and clear nail polish was applied around the coverslip perimeter.

Image stacks were acquired with 633 nm laser-line excitation using a Zeiss LSM 780 confocal microscope, at 20× or 40× magnification and 1024×1024 resolution centered around the patched soma, controlled by the ZEN2010 software (ZEISS International, Jena, Germany).

### Two-photon microscopy

2p laser-scanning microscopy was performed with a custom-built imaging workstation, as previously described (Abrahamsson et al., 2017). 2p excitation was achieved using a titanium-sapphire laser tuned to 820 nm for Alexa-594, and 930 nm for Alexa-488. Laser power output was monitored using a power meter and was controlled by adjusting a half-lambda plate passing the laser beam to a polarizing beam-splitting cube (Thorlabs GL10-B and AHWP05M-980). Laser gating was achieved with a mechanical shutter (Thorlabs SH05/SC10) triggered by software and scanning was achieved with 6215H 3-mm galvanometric mirrors (Cambridge Technology, Bedford, MA). Fluorescence was collected Semrock FF665 and laser light blocked with Semrock FF01-680. Red and green fluorescence were separated with Chroma t565lxpr combined with Chroma ET630/75M and Chroma ET525/50M. Bialkali photomultipliers (Scientifica 2PIMS-2000-20-20) were used in epifluorescence configuration to detect fluorescence. Laser-scanning Dodt contrast was achieved with custom optics, by collecting laser light that has passed through the acute slice with a spatial filter and a diffuser placed in a 1× telescope with an amplified diode (Thorlabs PDA100A-EC). Signals from photomultipliers and diode were acquired with a PCI-6110 digitization board (NI, Austin, TX, USA) using ScanImage 2019-2021 running in MATLAB (The MathWorks, Natick, MA, USA).

### Morphological reconstruction and analysis

After each whole-cell recording, cell morphologies were acquired at 40× as stacks of 512×512-pixel slices with 1 μm between each slice. Each slice is an average of 2 – 3 frames. These 3D stacks were Z-projected by maximum intensity, pseudo-colored, and assembled using Fiji (Schindelin et al., 2012).

Prior to reconstruction, brightness and contrast were adjusted in Fiji to ensure that branches in 3D stacks were optimally visible. Cells were then reconstructed by manual tracing using Neuromantic (Myatt et al., 2012). Reconstructed morphologies were analyzed with the qMorph in-house software, available at https://github.com/pj-sjostrom/qMorph (Zhou et al., 2021), and running in Igor Pro 9 (Wavemetrics Inc., Lake Oswego, OR, USA).

### Gap junction coupling quantification

To quantify gap junction coupling in astrocytes, individual cells were patched and filled with biocytin for 15 minutes. Biocytin-filled cells were fixed in PFA and stained, as described above. Z-stack images were acquired 25 μm above and 25 μm below the center of the patched cell soma with confocal imaging at 20× magnification as described above. Using the Cell Counter Plugin in Fiji, the number of fluorescent neighboring astrocyte soma was counted as a measure of amount of coupling.

### Imaging of spontaneous Ca^2+^ activity

Astrocytes were patched with glass pipettes loaded with Fluo-5F (200 μM) for 15 - 20 minutes. Using 2p excitation at 930 nm, the cell was imaged at a single focal plane for 180s, using 256×256-pixel frames acquired at 2.1 – 4.2 Hz frame rate. Ca^2+^ signals were measured as a change in green Fluo-5F fluorescence normalized to SR101 fluorescence acquired in the red channel.

At least 10 regions of interest (ROIs) of similar size were manually selected. We deliberately avoided automated ROI selection methods (e.g., Reynolds et al., 2017; Wang et al., 2019) because they are likely to affect signal decorrelation measurements, since automated signal-evolution-based ROI selection relies on pixel signal correlations. Baseline was set at the 10 – 40 frames with the weakest Ca^2+^ signal. To eliminate high-frequency noise, signal was low-pass filtered at 0.2 Hz. Mean Z-scores of Pearson’s r for the correlation of all ROIs were used to measure correlation of astrocyte Ca^2+^ activity.

Ca^2+^ events were detected by and counted with a simple thresholding algorithm, with the threshold detection level set to >1 sigma above background noise. Ca^2+^ event duration was similarly defined by the time during which the signal crossed this threshold. Frequency of Ca^2+^ activity was calculated by taking the total number of events detected divided by the total number ROIs selected for the cell and obtaining the number of events per ROI for each cell.

### Quantifying L5 cortical thickness

The thickness of cortical L5 was measured from confocal and 2p images using the brightfield and Dodt contrast channels, respectively. The upper and lower L5 boundaries were determined by visual inspection, with L5 pyramidal cells identified by their large soma and prominent apical dendrites. For each brain slices, the L5 thickness was measured in Fiji thrice and then averaged. Using Igor Pro, a sigmoid was numerically fitted to data, with sigmoid *rate* manually constrained.

### SR101 cell counts

Images were obtained from slices that were incubated in 1 μM SR101 for 5 minutes. Brightness and contrast were automatically set in FIJI, using histogram normalization. Cells were counted in one slice every ~3 - 10 μm in each stack. Dim cells or cells with incomplete soma were not counted. Cell counts for each stack were done at 3 different starting points and the average was taken to obtain cell density.

### Statistics

Unless otherwise noted, results are reported as the mean ± standard error of the mean (SEM). Significance levels are denoted using asterisks (* p < 0.05, ** p < 0.01, *** p < 0.001). Statistical tests were performed in Igor Pro. Pairwise comparisons were carried out using the two-tailed Student’s t-test for equal means. If the equality of variances F-test gave p < 0.05, we employed the unequal variances t-test. A t-test on Pearson’s r was used to determine significance of correlations. Sigmoids of the form 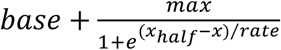 were numerically fit in Igor Pro.

## RESULTS

### Diverse astrocyte electrophysiology matured with age

To explore astrocyte electrophysiology within L5 visual cortex, we targeted astrocytes in P3 – P30 acute mouse brain slices for whole-cell patching. Since astrocytes can be difficult to identify using only brightfield or Dodt-contrast microscopy, we pre-incubated acute slices in 1 or 5 μM SR101 (see Methods), a fluorescent dye that is selectively taken up by astrocytes (**Fig. 1A**) (Nimmerjahn and Helmchen, 2012; Nimmerjahn *et al.*, 2004). With 2p imaging at 820 nm, we were able to visualize astrocyte staining in all cortical layers (**Fig 1B**), which enabled us to identify astrocytes for targeted patching under 2p microscopy (**Fig. 1C**).

**Figure 1.**
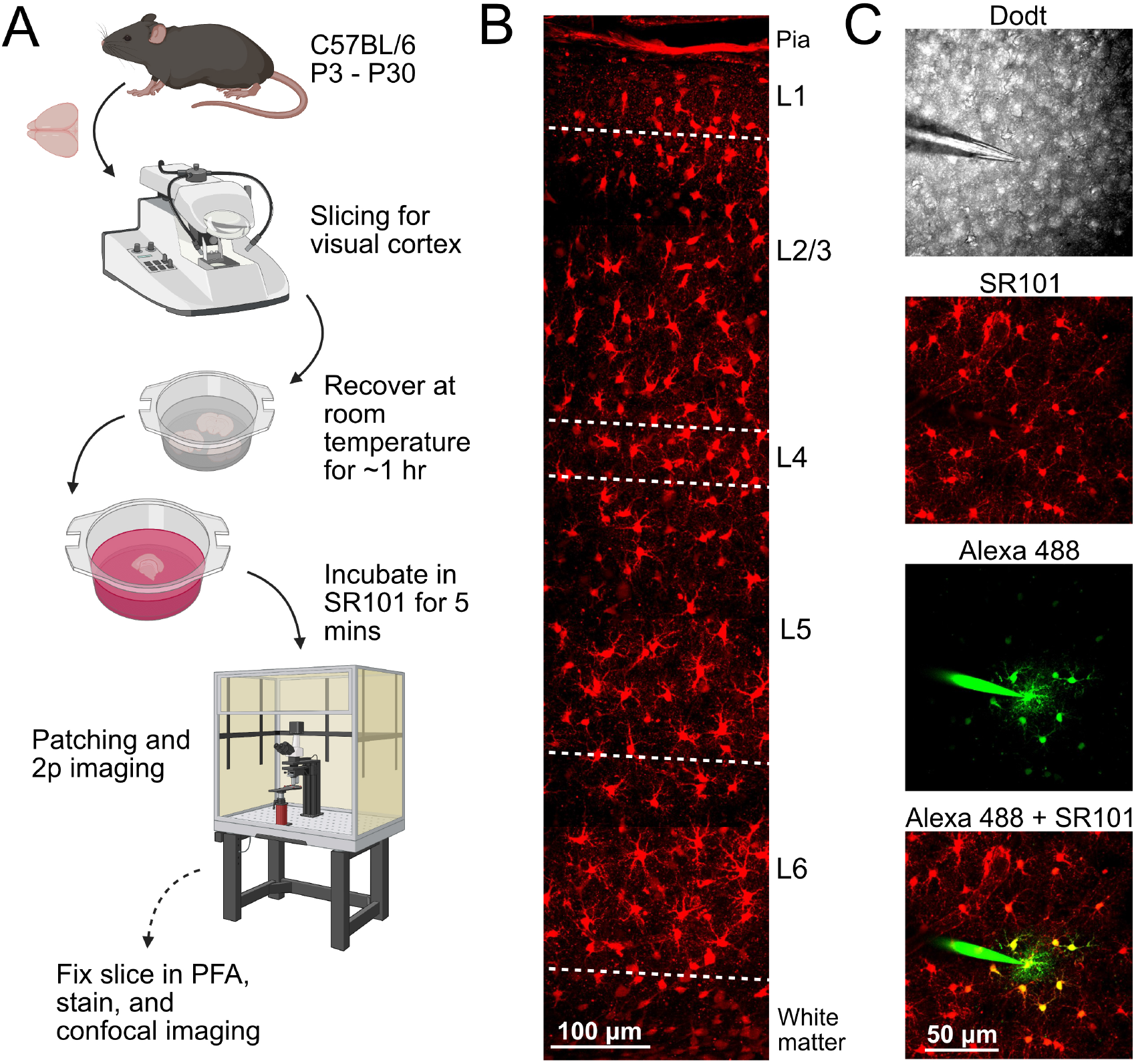
Astrocytes were targeted using SR101. (A) Acute slices were obtained from C57BL/6 mice and subsequently incubated in a container of ACSF to recover at room temperature for 1 hour. Prior to patching, a slice was transferred to a 6-well plate or petri dish filled with 1 or 5 μM SR101 for 5 minutes. A homemade mesh holder and bubbler allowed slices to sit on the mesh while the bubbler infused carbogen (not depicted) into the SR101 solution to keep the slice oxygenated. The slice was then transferred to the bath chamber of an electrophysiology rig for patching. After patching, the slice was either discarded or fixed in PFA for further analyses (optional dashed arrow). Schematic created in BioRender is for illustration purposes and does not show exact equipment used (see Methods). (B) Sample cortical stack of acute slice after pre-incubation in 1 μM SR101; Astrocytes in all cortical layers could be visualized under 2p microscopy at 930 nm. (C) SR101-labelled astrocytes (red) were targeted and patched with pipette filled with Alexa Fluor 488 (80 μM, green). The Alexa dye spread to neighboring astrocyte soma (yellow), presumably through gap-junctions.

Astrocyte identity was confirmed by verifying that the cell had characteristic passive properties (Adermark and Lovinger, 2008; Chai et al., 2017), including hyperpolarized Vm (−82 ± 0.3 mV, n = 127 cells, N = 43 animals) and low R_input_ (34 ± 2 MΩ). However, we found that R_input_ and V_m_ were variable, suggesting heterogeneity among astrocytes (**Fig. 2A**). Despite this heterogeneity, we could observe that astrocyte Vm increased with age, and that R_input_ decreased (**Fig. 2A**).

**Figure 2.**
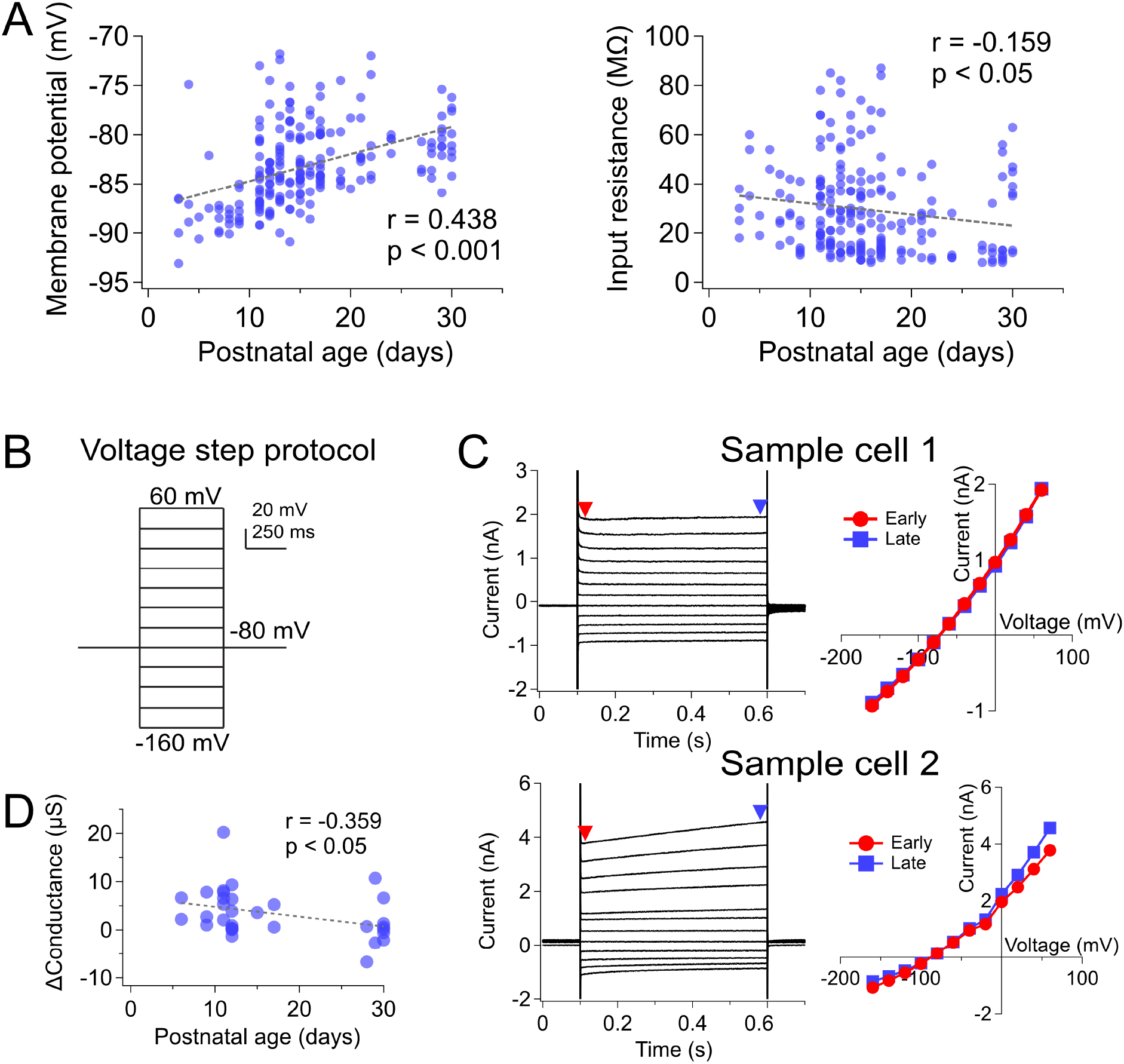
Membrane conductance properties were heterogeneous and matured with age. (A) Over development, resting Vm depolarized and R_input_ decreased. (B) To assess membrane biophysics, voltage steps of 20 mV decrements from +60 mV to −160 mV were applied to patched astrocytes. (C) Sample IV curves from two cells show different membrane conductance properties. Red arrowhead and round data points indicate early current readings (10 ms after start of voltage step) and, blue arrowhead and square data points indicate late current readings (10 ms before the end of voltage step). (D) The late-early conductance difference (see Methods) decreased as astrocytes matured (n = 30 cells, N = 12 animals).

To further investigate the membrane properties of astrocytes over development, we applied voltage steps from +60 mV to −160 mV to the patched cells (**Fig. 2B**). We saw cells exhibiting heterogenous membrane conductance properties, with some displaying passive current readings in response to the voltage steps (**Fig. 2C, top**), and others showing a time-dependent conductance, especially in the more depolarizing voltage steps (**Fig. 2C, bottom**). To quantify the time-dependent component, we took the difference in conductance between the late and early stages of the voltage step (see Methods) and compared them over development, which we found that this decreased over age (**Fig. 2D**).

### Gap junction formation may affect electrophysiological properties

We explored whether the changes observed in electrophysiological properties correlated with the development of gap-junctions in astrocytes. It is well known that astrocytes couple with neighboring astrocytes through connexin gap-junctions (Adermark and Lovinger, 2008; Schools et al., 2006). Therefore, we first patched individual astrocytes and dye-filled them with Alexa Fluor 488 for 30 minutes while taking time-lapse images every 2 minutes to visualize dye spreading to neighboring cells (**Fig. 3A, Supplemental video 1**). The rate of dye filling in astrocytes was quantified by measuring fluorescence intensity in a soma-centered ROI over time (**Fig. 3B**). The fluorescence intensity curves were fitted with a sigmoid to obtain their *x*_half_ values, i.e., the time at which cells were half filled. When these *x*_half_ values were independently clustered using a hierarchical agglomerative approach, two distinct groups emerged (purple, red, **Fig. 3C**). A positive correlation was also found between the *x*_half_ time and the distance between the neighbor and patched astrocyte (**Fig. 3D**). The two clustered groups likely correspond to primary neighbors that are coupled directly to the patched astrocyte (**Fig. 3 C, D**, purple and square) and secondary neighbors that are coupled to primary neighbors (**Fig. 3 C, D**, red and triangle). Finally, the *x*_half_ for secondary astrocytes (19 ± 0.4 min, n = 10) was larger than that of primary astrocytes (16 ± 0.5 min, n = 7; p < 0.001). These data show that primary neighbors closer to the patched astrocyte get dye-filled first, followed by secondary neighbors, on the timescale of minutes. Because Alexa 488 does not readily pass through connexin-30-containing gap junctions (Stephan et al., 2021), we opted to repeat the cell counts using biocytin. Using this method, we counted the number of dye-filled neighboring astrocytes (**Fig. 3E**) and found that this number increased with age (**Fig. 3F**). This suggested that astrocytes formed more gap junctions with their neighbors and/or that the gap junctions became more permeable over development (**Fig. 3G**). We also found that the increase in the number of coupled cells correlated with a decrease in R_input_, which itself decreased with age (**Fig. 3G, H**).

**Figure 3.**
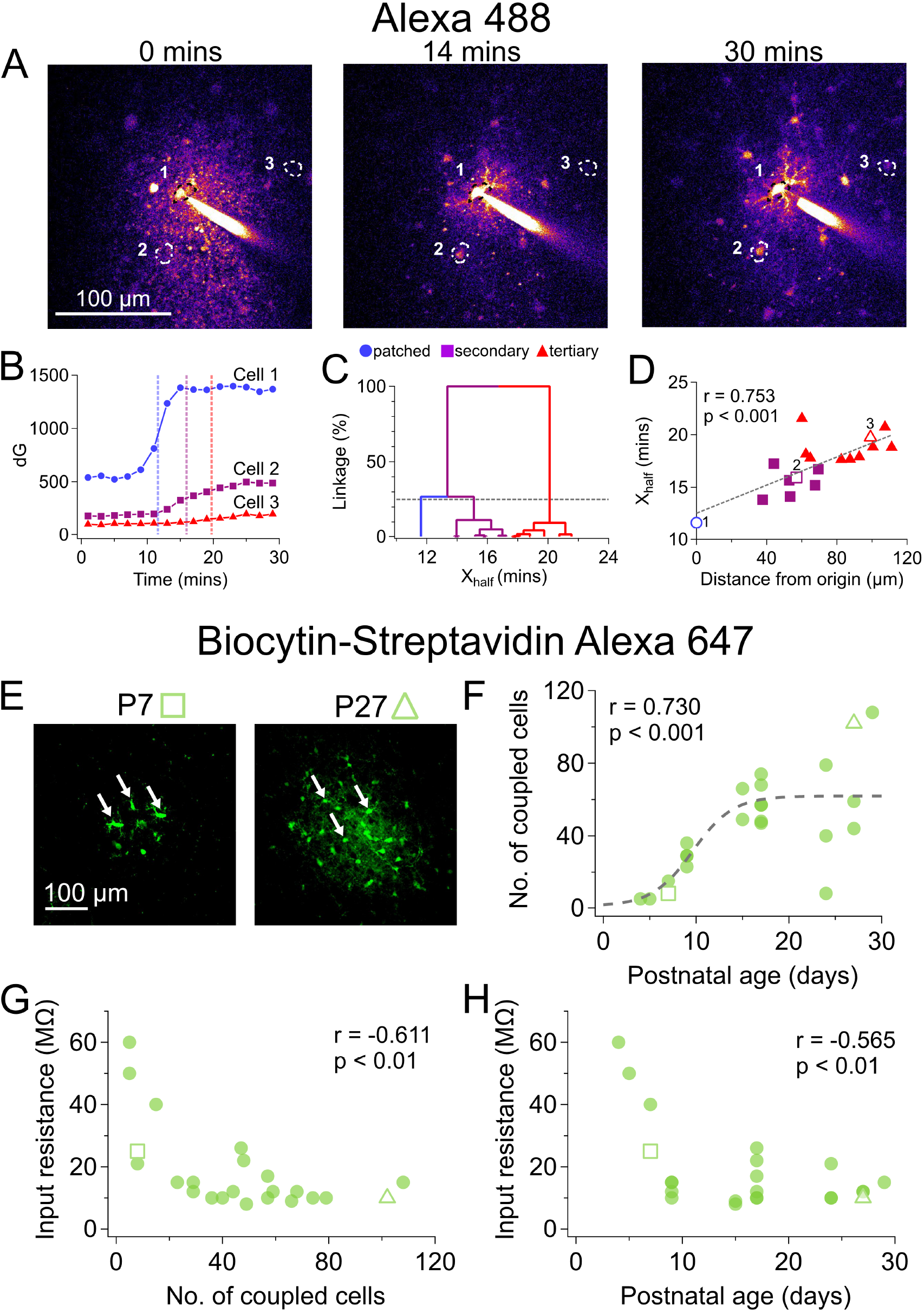
Astrocyte gap-junction coupling increased with age. (A) Sample 2p image shows Alexa-488 dye from patched astrocyte (Cell 1) spreading to example neighboring astrocytes Cell 2 and Cell 3 via gap junctions over time. (B) The fluorescence intensity signal of three sample astrocytes (1, 2, 3) in panel A increased. Fluorescence time courses and maxima were consistent with Cell 2 (purple) dye-filling directly from Cell 1 (blue), but Cell 3 (red) indirectly dye-filling from a secondary astrocyte such as Cell 2. Dashed lines denote the *x*_half_ half-max time points of sigmoidal fits (not shown). (C) The 25% best cut (dashed line) of hierarchically clustered *x*_half_ values independently suggested three fluorescence-onset clusters, consistent with the patched cell (blue) first dye-filling secondary astrocytes (purple), and those indirectly dye-filling tertiary cells (red) in a gap-junction-coupled astrocyte network. (D) Consistent with progressive dye-filling of a gap-junction-coupled astrocyte network, the fluorescence half-max *x*_half_ (see B, C) correlated positively with distance from patched astrocyte. Secondary (purple) and tertiary astrocyte labels (red) are based on clustering in C. Blue open circle is Cell 1 in A, purple open circle is Cell 2, and red open circle is Cell 3. (E) Since gap junctions may restrict passage of relatively large Alexa-488 dye molecules, we redid the experiments in A-D with biocytin, which is permeable through all astrocytic connexin channels (Stephan *et al.,* 2021). After histochemistry and confocal imaging (see Methods), stained cells (arrowheads) were counted. With this approach, sample P7 astrocyte did not dye-couple with neighboring cells as much as sample P27 astrocyte did. (F) The biocytin approach indicated that gap-junction coupling increased over development (n = 23 filled cells, N = 10 animals) and apparently plateaued around P15 but then remained quite heterogenous. Open square indicates P7 sample cell and open triangle indicates P27 sample cell depicted in (E). (G) & (H) R_input_ decreased with the number of coupled cells and with age, presumably related to gap junctions making astrocytes electrically leakier.

### Astrocyte arbors became denser, more compact, and more symmetric with age

To explore astrocyte morphology development, we manually reconstructed astrocytes at different ages from images obtained either with 2p or confocal imaging (**Fig. 4A**). We categorized cells obtained from animals into 3 age groups, those aged P1 – P10 (yellow), P11 – P20 (cyan), and P21 – P30 (magenta), for further comparison. To look at the extent of branching in these three groups, we compared the convex hulls, which measures the maximal reach of a cell’s processes. Surprisingly, although we expected the convex hulls to expand with age (i.e., that astrocytes became larger with age), they were mostly indistinguishable (**Fig. 4B**). However, when we looked at the branch density, we found that older astrocytes had more branches close to the soma as seen from a more saturated density heat map (**Fig. 4C**). This finding was supported by soma-centered Sholl analysis (Sholl, 1953), which showed a higher number of branch crossings close to the soma in old astrocytes (**Fig. 4D**). When we looked at the arbor center, i.e., the center of the entire reconstruction, and measured its distance from the soma, we found that astrocytes in the P1 – P10 and P11 – P20 groups had arbor centers further away from the soma compared to those in P21 – P30, meaning they were more asymmetric (**Fig. 4E**). When Sholl analysis for the three age groups were considered, we observed that overall, the cumulative number of crossings were greater in the older the age group (**Fig. 4F**). However, > 40 μm away from the soma, the older age groups stopped extending earlier in development (**Fig. 4F, inset**). Similarly, the cumulative crossings < 40 μm away from the soma showed a positive correlation with age (Fig. 4G), but not those > 40 μm away (**Fig. 4G, inset**).

**Figure 4.**
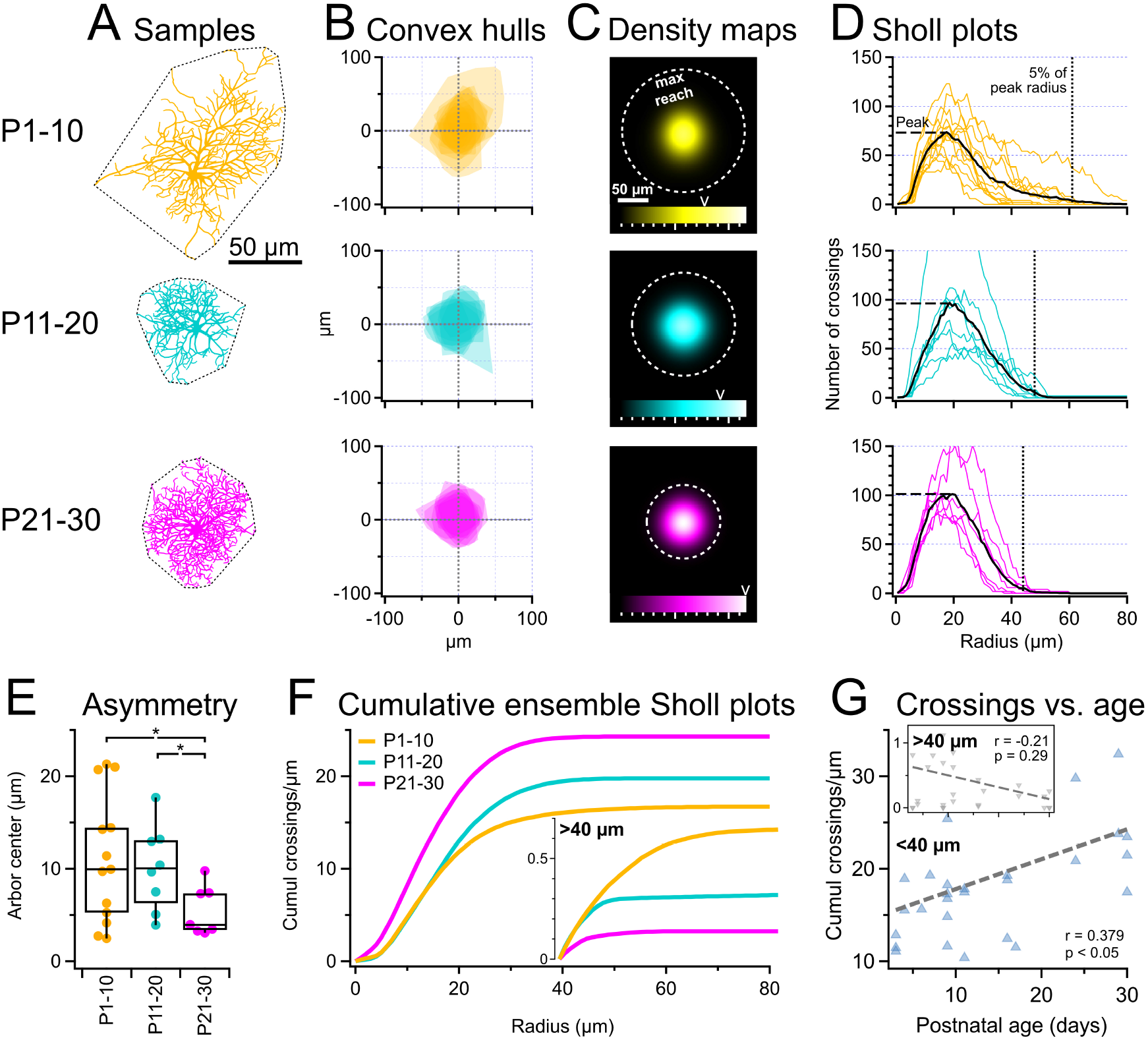
Maturing astrocyte arbors became denser, more compact, and more symmetric. (A) Sample astrocyte reconstructions illustrate how branching became denser, more compact, and more symmetric with age. The convex hull (dotted lines) indicates the maximal reach of branches. (B) Convex hulls, overlaid and centered on soma, suggested that with age, arbors became more symmetric as well as more compact. (C) Radially symmetric branch density heat maps (see Methods) suggested increased process branching close to the soma in old astrocytes. However, the farthest-reaching process seemed closer with age (dotted line). To enable comparison across age groups, heatmaps share scale, with map max intensity indicated by arrowhead above color scale. (D) In agreement with increased branching close to soma over age, Sholl analysis (Sholl, 1953) showed higher peak averages (horizontal dashed lines) and smaller 5%-of-peak-average radii (vertical dotted lines). Colored lines: Sholl plots of individual astrocyte arbors; black lines: age-group averages. (E) Distance from soma to arbor density center — a measure of arbor asymmetry — indicated that astrocyte arbors became more symmetric with development. (F) Cumulative ensemble Sholl plots across age groups indicated overall more branching in older astrocytes, although > 40 μm from the soma, young astrocytes branched more (inset). (G) In agreement with arborizations selectively enriching close to the soma, cumulative crossings within a 40 μm radius correlated positively with age but did not beyond 40 μm (inset; x-axis scale is identical).

### Astrocyte Ca^2+^ activity decorrelated with age

Astrocytes display spontaneous Ca^2+^ activity in their processes (Nett et al., 2002; Parri et al., 2001). We investigated if this Ca^2+^ activity changed with age. We targeted SR101-stained astrocytes P7 – P28 for patching and dialyzed them with the Ca^2+^-sensitive dye Fluo-5F (200 μM). We acquired 150-second-long videos of a single focal plane and selected ROIs to analyze fluorescent activity (**Fig. 5A**). To assess the similarity in Ca^2+^ activity between different ROIs, we calculated the Z-scored Pearson’s r pairwise across all ROIs in each recorded astrocyte. This way, we obtained a Ca^2+^ activity correlation matrix for each astrocyte (**Fig. 5B**). This heatmap was averaged to produce a mean Z-score for each astrocyte, which represents how correlated the Ca^2+^ activity was in that cell. By plotting astrocyte mean Z-score of Pearson’s r versus age (**Fig. 5C**), we found that Ca^2+^ activity decorrelated with age. In addition, we also detected and quantified the frequency and duration of individual Ca^2+^ transients and found that spontaneous Ca^2+^ signals became more frequent as well as shorter in duration with age (**Fig. 5C**). Taken together, these results indicate that early activity was dominated by single, large events that encompassed most or all branches in individual astrocytes (**Supplemental video 2**), whereas in more mature astrocytes, Ca^2+^ transients were more localized, relatively independent of each other, and comparatively brief (**Supplemental video 3**).

**Figure 5.**
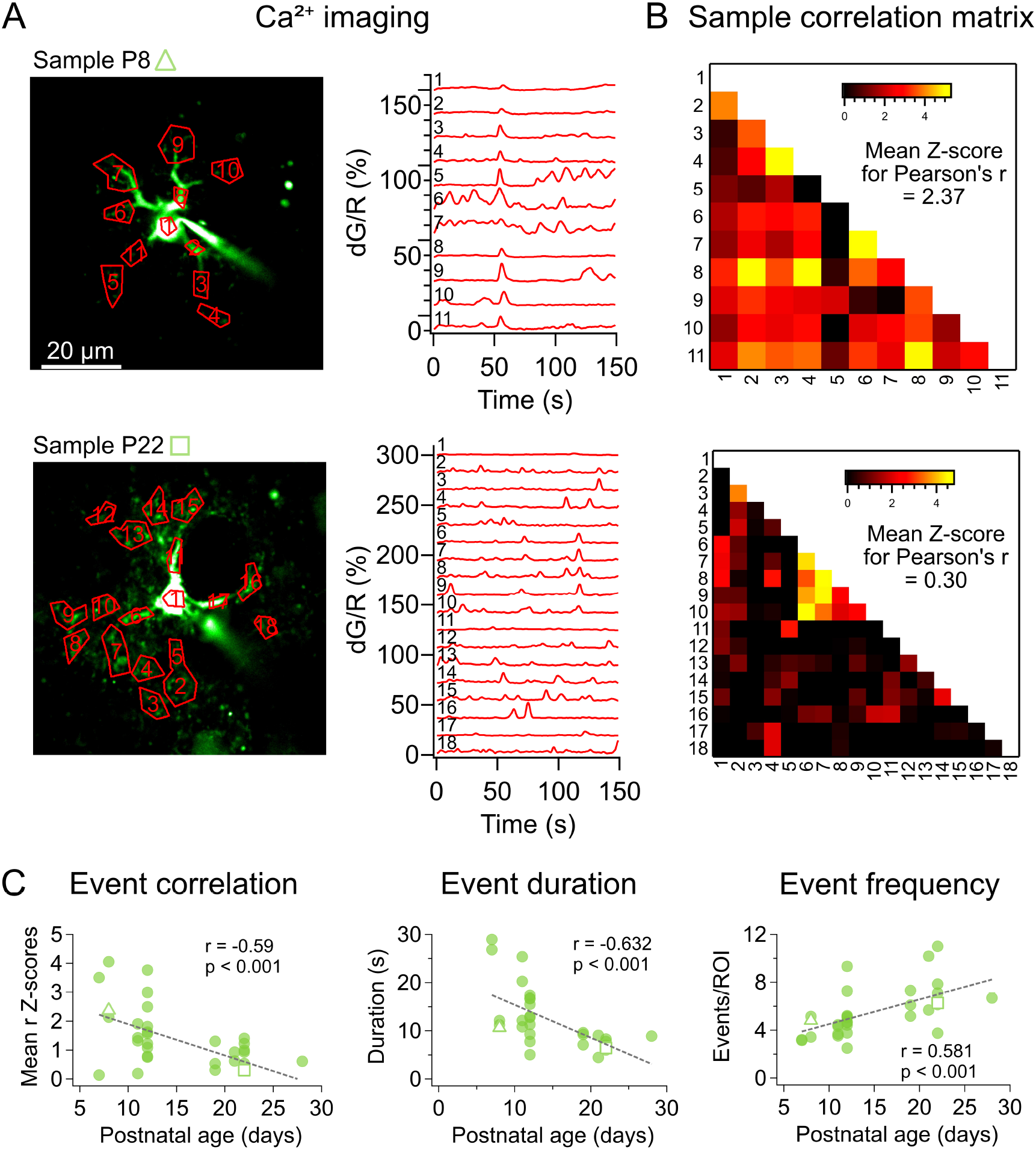
Ca^2+^ events became decorrelated, briefer, and more frequent with age. (A) Fluo-5F Ca^2+^ signals (green) were recorded as a 150-second-long movie (see Supplemental Movies 1 and 2) of these two sample P8 (open triangle) and P22 astrocytes (open square). During offline analysis, ROIs (left, red) were manually selected, and fluorescence was quantified as dG/dR sweeps (right). (B) Sample ROI cross-correlation matrices for the astrocytes in A. For each matrix, a single mean Z-score Pearson’s r was calculated, which was 2.3785 and 0.30477 for the P8 and P22 samples, respectively. Mean Z-score served as a metric for how correlated the activity of a cell was. (C) With age, astrocyte Ca^2+^ events became decorrelated, briefer, and more frequent (n = 31 cells, N = 12 animals). Open triangle indicated sample P8 cell and open square indicated sample P22 cell depicted in (A).

### Astrocyte density in L5 cortex remains constant while cortical thickness increases

We next measured L5 cortical thickness and found that it rapidly increases around P10, stabilizing at 250 μm thickness by ~P15 (**Supp. Fig. 1A**). Since our morphological reconstructions indicated that astrocytes did not grow larger with development (**Fig. 4**), we wondered if astrocytes therefore increased in numbers to ensure even tiling of the growing cortex. We therefore counted astrocytes in L5 and saw that their density remained relatively unchanged over age (**Supp. Fig. 1B**). Together with our morphological reconstructions (**Fig. 4**), these findings demonstrate that astrocytes grow in number rather than size. In summary, we found that L5 visual cortex astrocytes start off with few, probably partially overlapping branches, little gap-junction coupling, and correlated Ca^2+^ signaling, which mature by growing in numbers as L5 expands, by growing denser branches as tiling is established, by adding functional gap junctions, and by decorrelating Ca^2+^ signals into brief localized events (**Fig. 6**).

**Figure 6.**
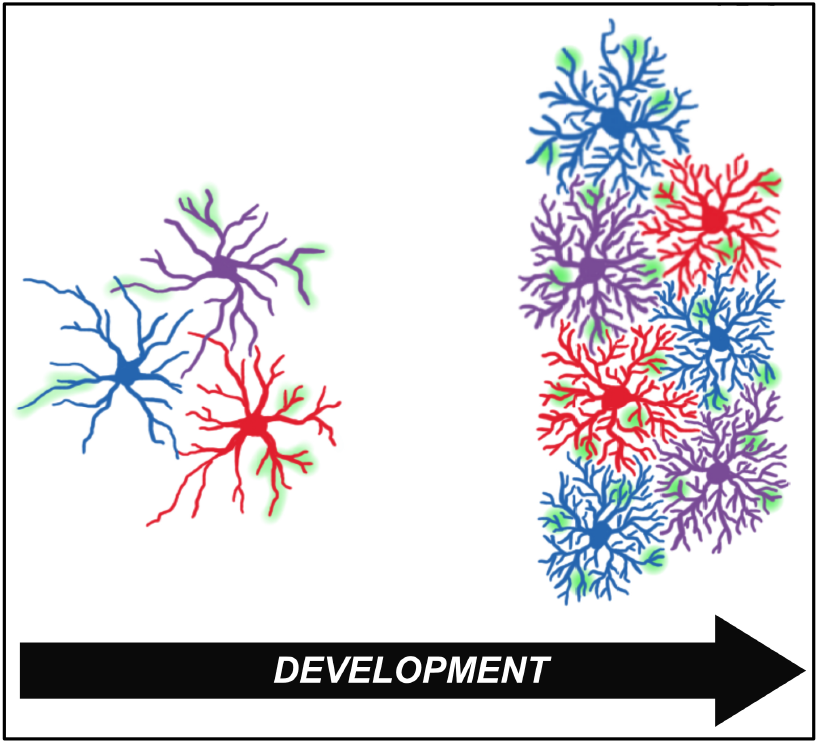
As astrocytes mature, branching, overall numbers, and Ca^2+^ signal compartmentalization increase. Proposed model illustrating key aspects of how young L5 V1 astrocytes — which are asymmetric, relatively far-reaching, and poorly branched (left) — mature by forming denser, more compact, and more symmetric arbors, as well as by growing more plentiful (right) to tile the growing neocortex. Ca^2+^ signals (green) also decorrelated and compartmentalized over development. Most properties stabilized by P15, right around eye opening, although arbors continued to refine by growing denser.

## DISCUSSION

Here, we studied the development of astrocytes in L5 of visual cortex across the ages P3 – P30. We found that astrocyte electrophysiological properties, although quite heterogeneous, mature by a depolarization of V_m_ and a reduction of R_input_. We also found that astrocyte morphology became denser and gap-junction coupling increased with development, and that spontaneous astrocyte Ca^2+^ activity decorrelated (**Fig. 6**). In addition, these Ca^2+^ transients increased in frequency and decreased in duration over the ages studied.

### Astrocyte arbors mature by growing denser and more compact

As expected, astrocytes developed denser arbors as they matured. In alignment with the findings of Bushong et al. (2004) our morphometry analysis revealed longer branches in the young compared to the old age group. Young arborizations were also more asymmetric. This suggests that astrocytes first grow long but weakly branched arbors that are subsequently pruned while arborizations grow dense close to the soma. In this view, astrocyte branches may dynamically develop by growth and retraction, perhaps to establish even tiling by sampling different territories.

Astrocyte arbors may thus develop like neuronal axons and dendrites do, by initial exuberance followed by subsequent pruning (Riccomagno and Kolodkin, 2015). As tiling boundaries form, the territory covered fills in with additional branches as the astrocyte matures. However, more detailed study is required to verify this growth-and-retraction idea, e.g., by in-vivo time lapse imaging. Nevertheless, our findings are consistent with e.g., the report of Bushong *et al.* (2004) who found heterogenous astrocyte morphologies in rat hippocampus before P14. They also found that astrocytes extended long processes and that astrocyte tiling was not consistently seen until P14 (Bushong *et al.,* 2004).

With SR101 staining, we showed that astrocyte density remains relatively stable between P3 – P30, which means astrocyte spacing is constant. Therefore, the longer branching of P1 – P10 astrocytes suggests that they might extend into the domains of their neighbors. During maturation, branches may therefore retract to enable the eventual astrocyte tiling in the mature brain. However, additional work is required to put this idea to the test.

### Astrocytes depolarize and become leaky as they mature

Astrocyte biophysics was heterogenous across cells, with for example R_input_ spanning an order of magnitude, and with some but not all astrocytes carrying a slowly adapting conductance. Despite this variability, we found developmental changes: With age, astrocytes depolarized and became leakier, while the slowly adapting current gradually decreased (**Fig. 2**). At ~P15, membrane properties seemed to stabilize.

The heterogenous electrophysiology we report has previously been found in astrocytes of striatum, hippocampus, and neocortex (Adermark and Lovinger, 2008; McKhann et al., 1997). It has been argued that such heterogeneity reflects differences in gap-junction coupling (McKhann et al., 1997) or astrocyte types (Adermark and Lovinger, 2008). Anders et al. (2014) showed that, in the hippocampus, gap-junction coupling relates to astrocyte location relative to the pyramidal cell layer, suggesting the existence of layer-dependent astrocyte properties. We, however, only studied astrocytes in L5 of visual cortex, suggesting that the heterogeneity we found is not due to layer location, but more likely gap-junction coupling (McKhann *et al.,* 1997).

Astrocyte gap-junction coupling first develops in rat visual cortex at P1 (Schools et al., 2006) and then gradually increases (Stephan et al., 2021). In mouse thalamus, Griemsmann et al. (2015) found that astrocyte coupling did not increase after the first two postnatal weeks. In agreement, we found that dye-coupling was stabilized from P15 and onwards (**Fig. 3**). This may suggest that developmental gap-junction coupling is linked to the maturation of membrane properties, due to a correlation of these properties (**Fig. 3G, H**), which is also supported by prior experiments done in the striatum (Adermark and Lovinger, 2008).

### Astrocytes Ca^2+^ events mature by decorrelating and shortening

Spontaneous Ca^2+^ activity in astrocyte processes have been linked to developmental circuit plasticity (Aguado *et al.*, 2002; Fiacco and McCarthy, 2004; Parri *et al.,* 2001). We showed how spontaneous Ca^2+^ events developed alongside astrocyte electrophysiology and morphology, stabilizing from ~P15 onwards with relatively short, decorrelated, and compartmentalized Ca^2+^ transients. A previous theoretical study suggested that morphological profile determines frequency of spontaneous astrocyte Ca^2+^ signals (Wu et al., 2019), specifically that spontaneous Ca^2+^ events start in thin astrocytic processes due to their high surface area-to-volume ratio, which promotes Ca^2+^-induced Ca^2+^ release. In agreement, computational modelling has reproduced spontaneous Ca^2+^ signals by Ca^2+^-induced Ca^2+^ release in fine astrocyte processes (Denizot et al., 2019).

### Conclusions and future directions

Our present work provides a descriptive foundation of astrocyte maturation in juvenile mouse visual cortex. Since astrocyte Ca^2+^ signaling is key to neocortical plasticity (Min and Nevian, 2012), our findings outline a framework for the study of astrocyte-mediated control of visual cortex critical period plasticity, e.g., by showing that biophysics and Ca^2+^ signaling of visual cortex astrocytes were relatively mature by eye opening.

As astrocyte branching grew denser, Ca^2+^ signals decorrelated. This was not surprising, because finer arborization should naturally dissociate Ca^2+^ events as branches compartmentalize. Astrocyte morphology, however, continued to mature after P15 when biophysics and dye-coupling were stabilized. Whether maturation of morphology determines developmental decorrelation of astrocyte Ca^2+^ activity thus remains unclear in our hands.

Our study suggests that — like axonal and dendritic arbors (Riccomagno and Kolodkin, 2015) — astrocyte branches may initially extend too far and later be pruned to establish tiling (**Fig. 6**). In the mammalian central nervous system, it is known that astrocytes and microglia participate in axon pruning (Riccomagno and Kolodkin, 2015). If astrocytes themselves are pruned, this raises the intriguing question: who prunes the pruners?

## LIMITATIONS OF THE STUDY

A potential caveat is that we measured astrocyte asymmetry in 2D. A 3D analysis might provide more context to the positioning of astrocytes relative to other local structures. For example, Refaeli et al. (2021) showed that astrocytes display orientation preference in hippocampus relative to the CA1 pyramidal area. However, a more refined 3D analysis should not alter our overall finding that arborizations mature by growing denser, more compact, and more symmetric.

Another potential caveat is that we pooled morphological data obtained with 2p microscopy of cells dye-loaded during patch-clamp recording with data obtained by confocal imaging after biocytin histology. However, we previously found these methods offered indistinguishable morphological cell classification performance (Blackman et al., 2014), which justified pooling these types of data. In agreement, each data set alone offered similar outcome (not shown).

The finest astrocyte compartments cannot be resolved with confocal or 2p microscopy but require super-resolution or electron microscopy (Arizono et al., 2022). Confocal and 2p imaging may thus underestimate branch numbers. However, we had no problems detecting morphological changes across P3 – P30, suggesting a sufficiently detailed resolution for our purposes.

SR101 labels only a subset of astrocytes, although the developmental profile of SR101-positive cells is comparable to that of other astrocytes (Kafitz et al., 2008). Additionally, SR101 has limited specificity and may also label oligodendrocytes, although that requires longer incubation time than we used here (Hill and Grutzendler, 2014). Oligodendrocyte morphology is furthermore quite distinct from that of astrocytes.

Finally, in several cases, causation remains to be established. For example, the relationship between gap-junction coupling and R_input_ was correlational, not causal.

## Supporting information

Supplemental Video 1 | Sample time lapse movie of Alexa 488 filling an astrocyte.

Supplemental Video 2 | Sample Ca2+ activity in P8 astrocyte.

Supplemental Video 3 | Sample Ca2+ activity in P22 astrocyte.

## ACKNOWLEDGEMENTS

We thank Alanna Watt, Hovy Wong, Christina Chou, Sabine Rannio, Amanda McFarlan, Shawniya Alageswaran, and members of the Sjöström lab for help and useful discussions. The schematic in Figure 1A was created with BioRender.com.

## AUTHOR CONTRIBUTION

Conceptualization, AW and PJS; Methodology, AW, CG, and PJS; Software, PJS; Validation, AW and PJS; Formal Analysis, AW and PJS; Investigation, AW and CG; Resources, PJS; Data Curation, AW, CG, and PJS; Writing – Original Draft, AW and PJS; Writing – Reviewing and Editing, AW and PJS; Visualization, AW and PJS; Supervision, AW and PJS; Project Administration, AW and PJS; Funding Acquisition, AW and PJS.

## DATA AVAILABILITY STATEMENT

The raw data supporting the conclusions of this manuscript will be made available by the authors, without undue reservation, to any qualified researcher.

## SUPPLEMENTARY DATA

### Sample Ca^2+^ movies

***Supplemental Video 1 | Sample time lapse movie of Alexa 488 filling an astrocyte.***

***Supplemental Video 2 | Sample Ca2+ activity in P8 astrocyte.***

***Supplemental Video 3 | Sample Ca2+ activity in P22 astrocyte.***

### SR101+ astrocyte density remains stable over age

**Supplemental Figure 1.**
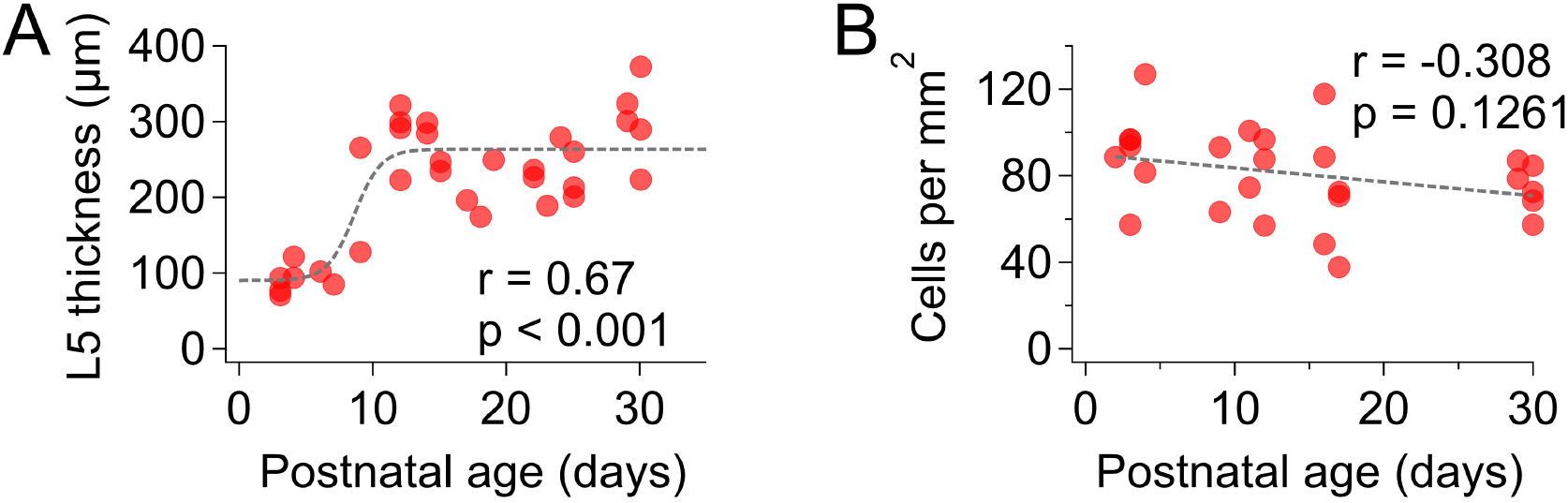
L5 cortical thickness increased but astrocyte density in L5 remains stable over age. (A) L5 cortical thickness (see Methods) roughly tripled over the examined age range (n = 32 slices, N = 24 animals). (B) Yet, astrocyte cell density remained relatively unchanged over this age span (n = 26 images, N = 13 animals).

